# Machine Learning Models to Interrogate Proteomewide Covalent Ligandabilities Directed at Cysteines

**DOI:** 10.1101/2023.08.17.553742

**Authors:** Ruibin Liu, Joseph Clayton, Mingzhe Shen, Shubham Bhatnagar, Jana Shen

**Affiliations:** Department of Pharmaceutical Sciences, University of Maryland School of Pharmacy, Baltimore, MD 21201, USA; Division of Applied Regulatory Science, Office of Clinical Pharmacology, Center for Drug Evaluation and Research, U.S. Food and Drug Administration, Silver Spring, MD 20993, USA; Department of Computer Science, University of Maryland at College Park, College Park, MD 20742, USA

**Keywords:** Covalent drug discovery, machine learning, protein structures, database, AlphaFold, kinases

## Abstract

Machine learning (ML) identification of covalently ligandable sites may accelerate targeted covalent inhibitor design and help expand the druggable proteome space. Here we report the rigorous development and validation of the tree-based models and convolutional neural networks (CNNs) trained on a newly curated database (LigCys3D) of over 1,000 liganded cysteines in nearly 800 proteins represented by over 10,000 three-dimensional structures in the protein data bank. The unseen tests yielded 94% and 93% AUCs (area under the receiver operating characteristic curve) for the tree models and CNNs, respectively. Based on the AlphaFold2 predicted structures, the ML models recapitulated the newly liganded cysteines in the PDB with over 90% recall values. To assist the community of covalent drug discoveries, we report the predicted ligandable cysteines in 392 human kinases and their locations in the sequence-aligned kinase structure including the PH and SH2 domains. Furthermore, we disseminate a searchable online database LigCys3D (https://ligcys.computchem.org/) and a web prediction server DeepCys (https://deepcys.computchem.org/), both of which will be continuously updated and improved by including newly published experimental data. The present work represents a first step towards the ML-led integration of big genome data and structure models to annotate the human proteome space for the next-generation covalent drug discoveries.

## INTRODUCTION

Over the past two decades, targeted covalent inhibitor (TCI) discovery has become mainstream in the efforts to overcome limitations of traditional reversible inhibitors and expand the druggable proteome space. ^1–3^ In the TCI design, an electrophilic functional group (also known as the warhead) is incorporated into a reversible ligand to enhance potency, selectivity, and target residence time or to inhibit a previously deemed undruggable protein, e.g., KRAS-G12C that lacks of a traditional ligandable pocket for reversible binding. ^4^ An irreversible and sometimes also reversible covalent bond is formed between the warhead and a nucleophilic (reactive) amino acid residue in the target protein. Cysteine is the most nucleophilic amino acid and has been most popular for covalent ligation. In fact, all FDA approved TCIs are directed at a cysteine. ^5^

In silico approaches hold a great promise to accelerate the proteome-wide TCI discovery efforts. In recent years, molecular dynamics (MD) simulations ^6–8^ have been developed to assess cysteine reactivities and ligandabilities; however, they cannot be scaled up to the proteome level due to the high computational cost. Machine learning (ML) models trained on the cysteine-liganded co-crystal structures in the protein data bank (PDB) have also been reported to evaluate the cysteine ligandabilities. In the earlier work, the support vector machine (SVM) models were trained on 1057 cysteine-liganded co-crystal structures of 515 proteins and achieved the best AUC (area under the curve of receiver operating characteristic or ROC), recall, and precision of 0.73, 0.62, and 0.41, respectively, ^9^ in an unseen test. In a most recent publication, ^10^ the graph neural network (GNN) models DeepCoSI were trained on the CovalentInDB database which contains 1,042 cysteine-liganded co-crystal structures of 259 proteins, with the best training AUC of 0.92.

The emergence of the powerful and continuously improving AlphaFold2 (AF2) structure prediction engine ^12^ further underscores the potential utility of structure-based ligandability prediction tools. In this work, we present a new database LigCys3D (https://db.computchem.org/), which annotates 1,133 ligandable cysteines of 778 proteins and their X-ray crystal structure representations in the PDB, including the cysteine-liganded and unliganded forms. Employing this database, we trained and validated two types of ML models, the decision tree models and the three-dimensional convolutional neural networks (3D-CNNs). To our best knowledge, LigCys3D is the largest to date and significantly surpasses those used for the previous ML models ^9,10^ in terms of both the number of unique cysteines, proteins, as well as the number of structural representations. Another uniqueness of this work is the inclusion of ligand-free (apo) X-ray structures in the training set, which is expected to increase the extrapolation power of the ML models. Finally, development of decision tree models (as opposed to the black box neural networks) based on physicochemical features allows us to dissect the physical determinants and gain systematic understanding of covalent ligandabilities. In multiple unseen tests, the XGBoost models and CNNs delivered the AUCs of 94%±1% and 93%±3%, respectively. The models were also applied to recapitulate the newly liganded cysteines based on AF2 predicted structures, giving recall values over 90%. Finally, a web server https://deepcys.computchem.org/ was developed to assist the community of covalent drug discovery.

## RESULTS and DISCUSSION

### Construction of a structure database of cysteine-liganded proteins determined by crystallography

In order to train ML models, we first built a database of proteins containing cysteines that have been covalently modified by ligands. The recently published CovPDB ^13^ and CovalentInDB ^10^ databases together contain 659 liganded cysteines in 484 unique proteins. We performed an exhaustive search in the PDB and found additional 474 liganded cysteines in 296 unique proteins. Together, we compiled 1,133 liganded cysteines in 778 unique proteins. These cysteines will be referred to as positives. The rest of the 3,077 cysteines in these proteins are unliganded, which will be referred to as negatives. We note, although the unliganded cysteines are more reliable negatives than the cysteines in proteins that have not been cysteine-liganded before, false negatives are still possible. Using the most recent PDBx/mmCIF files by SIFTS, ^14^ we matched each cysteine with the (gene) accession number and residue ID in the UniProt knowledge base (UniProtKB). ^11^ 76% of the cysteine-liganded proteins are enzymes, including 101 proteases, 59 kinases and 433 other enzymes (Fig. 1a). Channels/transporters/receptors (58), transcription factors and regulators (41) are also present, along with 66 proteins that do not have functional classifications based on UniProtKB ^11^ or SCOP2^15^(Fig. 1a).

**Figure 1.**
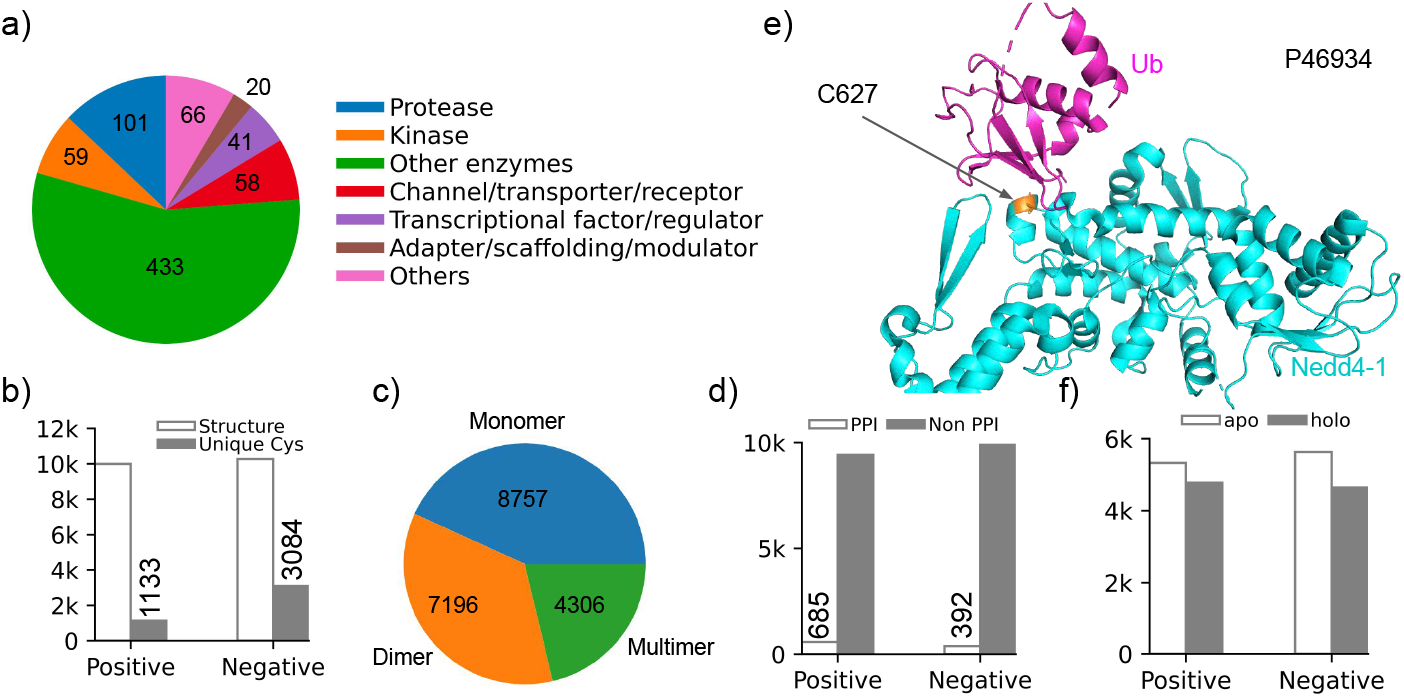
Analysis of the ligandable cysteines and the associated X-ray structures in the down-sampled LigCys3D. **a)** Functional classes of the proteins that have at least one ligandable (positive) cysteine according to the structures deposited in the PDB. Functional information is taken from the UniProtKB. ^11^ **b)** Number of positive and negative cysteines, and the number of PDB structures representing these cysteines. A positive cysteine is represented by up to 10 and an average of ∼9 structures, while a negative cysteine is represented by up to 4 and an average of ∼3 structures. **c)** Number of nonunique cysteines that are in monomer, dimer, and multimer structures based on the biological assembly information in the PDB. **d)** Number of PDB structures that represent positive or negative cysteines that are near the PPI or not. A PPI cysteine was defined using a distance cutoff of 4.5 Å between the sulfhydrl sulfur and the nearest heavy atom in another chain. **e)** Nedd4-1 (cyan) contains a cysteine (C627, orange) at the PPI with ubiquitin (magenta) in the PDB entry 5C7J. While not liganded in this structure, Cys627 is liganded by a covalent inhibitor in a different, monomeric structure (PDB ID: 5C91). **f)** Number of PDB structures that are apo (ligand free) or holo (bound to any ligand) for positive and negative cysteines.

The CovPDB ^13^ and CovalentInDB ^10^ databases contain only the cysteine-liganded PDB structures, based on which the previously reported ML models were trained. ^9,10^ This is not ideal, as the conformational variability is neglected, which may limit the model transferability (see later discussion). Thus, we augmented the dataset to a total of 10,105 positive entries (10,105 X-ray structures representing 1,133 positive cysteines) and 97,754 negative entries (97,754 X-ray structures representing 3,084 unique negative cysteines). The quaternary structure was built based on the bioassembly information in the PDB. On average, each positive cysteine is represented by ∼9 structures, and in most of these structures the positive cysteine is not liganded, i.e., the structure is either ligand free or in complex with a reversible ligand. We will refer to this dataset as LigCys3D (ligandable cysteine three-dimensional structure database) hereafter. Since there are significantly more negatives than positives, we randomly down-sampled the negative entries to 10,267, i.e., 10,267 X-ray structures representing 3,084 negative cysteines (an average of ∼3 structures per negative cysteine). In total, 20,259 entries were curated as the dataset for model hold-out, training and cross validation (CV, Fig. 1b).

### Structural diversity, variability, and allostery are represented in the augmented dataset

Considering the quaternary structures associated with the entries, 8,757 are monomers, 7,196 are dimers, and 4,306 multimers (Fig. 1c). In addition, 685 structures associated with the (119) positive cysteines and 392 structures associated with the (110) negative cysteines are located at the protein-protein (or protein-nucleic acid) interfaces (PPIs, Fig. 1d), as defined using a distance cutoff of 4.5 Å between the cysteine sulfur and its nearest heavy atom from a different chain in the PDB file. An interesting PPI example is the HECT E3 ubiquitin ligase Nedd4-1, which regulates metabolism, growth, and development and is a promising target for treating cancers and other diseases (Fig. 1e). ^16^ Nedd4-1 has a noncatalytic cysteine C627, which is located at the binding interface with ubiquitin (PDB ID: 5C7J) ^17^ and has been modified by a covalent inhibitor (PDB ID: 5C91). ^16^ In addition to the cysteine-liganded structures, through data augmentation the positive cysteines are also represented by co-crystal structures in complex with reversible ligands as well as surprisingly more than 50% ligand-free structures. For the positive entries, 3,912 are ligand free and 3,695 are ligand bound, while for the negative entries, 3,601 are ligand free and 3,003 are ligand bound (Fig. 1f). These analyses demonstrate that our data augmentation strategy affords structure diversity and variability, which we surmised to be essential for training truly predictive and transferable models. The inclusion of structural variation may also help with the detection of cryptic pockets. ^18^ We should also note that in the LigCys3D dataset, each protein has on average 1.5 ligandable cysteines, which suggests that allosteric sites are also represented.

### The top three tree models are highly predictive of lig-andable cysteines

The recent constant pH MD titration simulations of a large number of kinases uncovered common structural and physical features for reactive cysteines (high tendency to deprotonate at physiological pH) and ligandable cysteines. ^6,8,19,20^ Thus, we surmised that the feature-based ML classification models such as decision trees may be suited for predicting cysteine ligandabilities. Based on the findings from these studies ^6,8,19,20^ we devised a set of descriptors (37 after removal of multicollinearity, see Methods) for training the tree-based classifiers using PyCaret. ^21^ Given the small training dataset, tree models avoid the over-fitting problem that plagues the more sophisticated models that make use of vast parameter space. From the down-sampled LigCys3D, 10% of the entries were randomly picked as hold-out for the “unseen” test, while the remaining 90% of the entries were reserved for training/CV. UniProt accession number and residue IDs were used to ensure cysteines are unique between the training/CV and test sets. The 10-fold CV was used, where different folds have unique cysteines. Following the CV, the model was retrained with hyperparameter tuning before being applied to the test set. This process (data splitting, training/CV, and test) was repeated six times to generate statistics for model evaluation. To verify that the data splitting is unbiased and to generate more robust statistics, we also repeated the above process 30 times for the ET model. The resulting metrics are very similar, with the test AUC value unchanged (Table S1).

The eXtreme Gradient Boosting (XGBoost), Extra Tree (ET), and Light Gradient Boosting (LightGBM) are the top three best performing models according to the AUC, recall, precision, and F1 score in the unseen tests (Table 1). These four metrics analyze the model performance in different ways. The AUC is an aggregate measure of true and false positive rates across all possible classification thresholds. Recall measures the accuracy of the positive predictions given a threshold (percentage of the predicted positives that are truly positive), while precision measures the percentage of positive entries correctly identified. The F1 score is the harmonic mean of recall and precision. Note, we also calculated the selectivity and negative predictive value (NPV), which respectively measure the accuracy and precision of predicting negatives. These metrics are de-emphasized in this work because our training set might contain false negatives as discussed before and knowing the positives are more relevant in drug discovery.

**Table 1.**
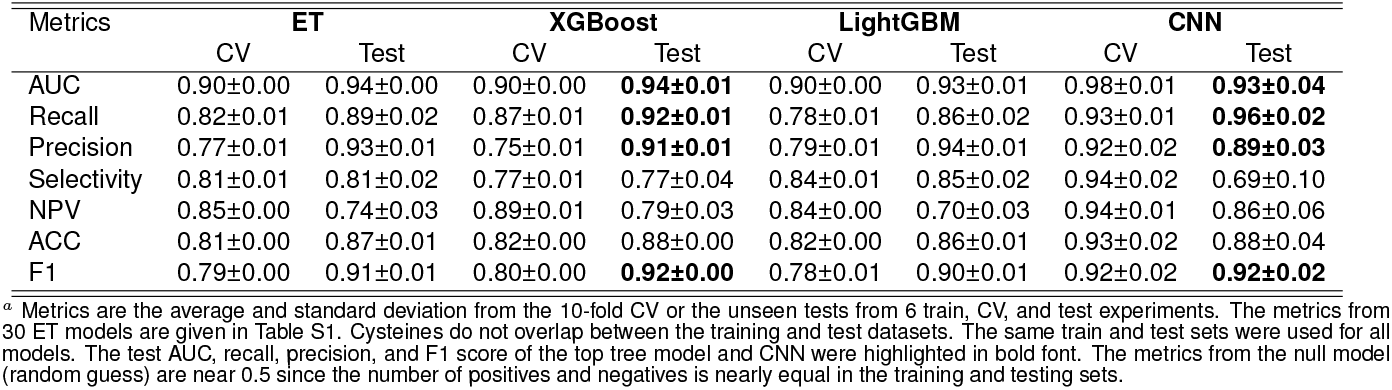
Performance of the tree-based and CNN models in the cross validations and unseen tests^*a*^.

| Metrics | ET |  | XGBoost |  | LightGBM |  | CNN |  |
| --- | --- | --- | --- | --- | --- | --- | --- | --- |
|  | CV | Test | CV | Test | CV | Test | CV | Test |
| AUC | 0.90±0.00 | 0.94±0.00 | 0.90±0.00 | **0.94±0.01** | 0.90±0.00 | 0.93±0.01 | 0.98±0.01 | **0.93±0.04** |
| Recall | 0.82±0.01 | 0.89±0.02 | 0.87±0.01 | **0.92±0.01** | 0.78±0.01 | 0.86±0.02 | 0.93±0.01 | **0.96±0.02** |
| Precision | 0.77±0.01 | 0.93±0.01 | 0.75±0.01 | **0.91±0.01** | 0.79±0.01 | 0.94±0.01 | 0.92±0.02 | **0.89±0.03** |
| Selectivity | 0.81±0.01 | 0.81±0.02 | 0.77±0.01 | 0.77±0.04 | 0.84±0.01 | 0.85±0.02 | 0.94±0.02 | 0.69±0.10 |
| NPV | 0.85±0.00 | 0.74±0.03 | 0.89±0.01 | 0.79±0.03 | 0.84±0.00 | 0.70±0.03 | 0.94±0.01 | 0.86±0.06 |
| ACC | 0.81±0.00 | 0.87±0.01 | 0.82±0.00 | 0.88±0.00 | 0.82±0.00 | 0.86±0.01 | 0.93±0.02 | 0.88±0.04 |
| F1 | 0.79±0.00 | 0.91±0.01 | 0.80±0.00 | **0.92±0.00** | 0.78±0.01 | 0.90±0.01 | 0.92±0.02 | **0.92±0.02** |

<sup>a</sup> Metrics are the average and standard deviation from the 10-fold CV or the unseen tests from 6 train, CV, and test experiments. The metrics from 30 ET models are given in Table S1. Cysteines do not overlap between the training and test datasets. The same train and test sets were used for all models. The test AUC, recall, precision, and F1 score of the top tree model and CNN were highlighted in bold font. The metrics from the null model (random guess) are near 0.5 since the number of positives and negatives is nearly equal in the training and testing sets.

**Table 2.**
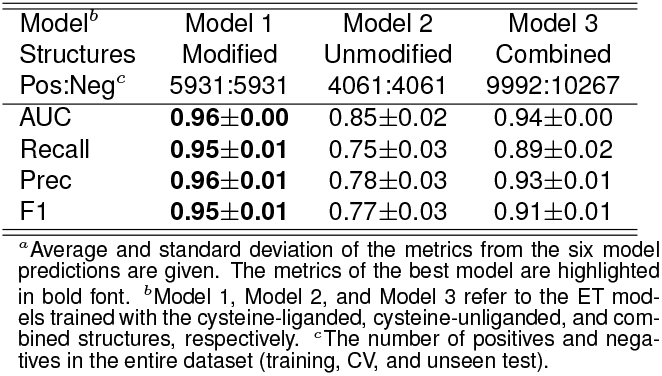
Impact of training with unmodified structures on the ET model predictions^*a*^.

The best model XGBoost gave an AUC of 0.94±0.01 (Fig. 2b) and a maximum F1 score of 0.92±0.02, which was achieved at the threshold value of 0.30 (Fig. 2c). With this threshold, the recall and precision are 0.92±0.01 and 0.91±0.01, respectively (Table 1). The test metrics of the ET classifier closely follow those of the XGBoost. Considering the test AUC, recall, and precision of 0.93–0.94, 0.89–0.96, and 0.89–0.91, respectively, the top three tree-based models are highly predictive of the ligandable cysteines. Note, some test metrics of the tree models are higher than those of the CVs. This is because the CV metrics were calculated by averaging the models trained on 9 folds of data, while the test metrics were for the model trained on the entire 10 fold of data with the optimized hyperparameters.

**Figure 2.**
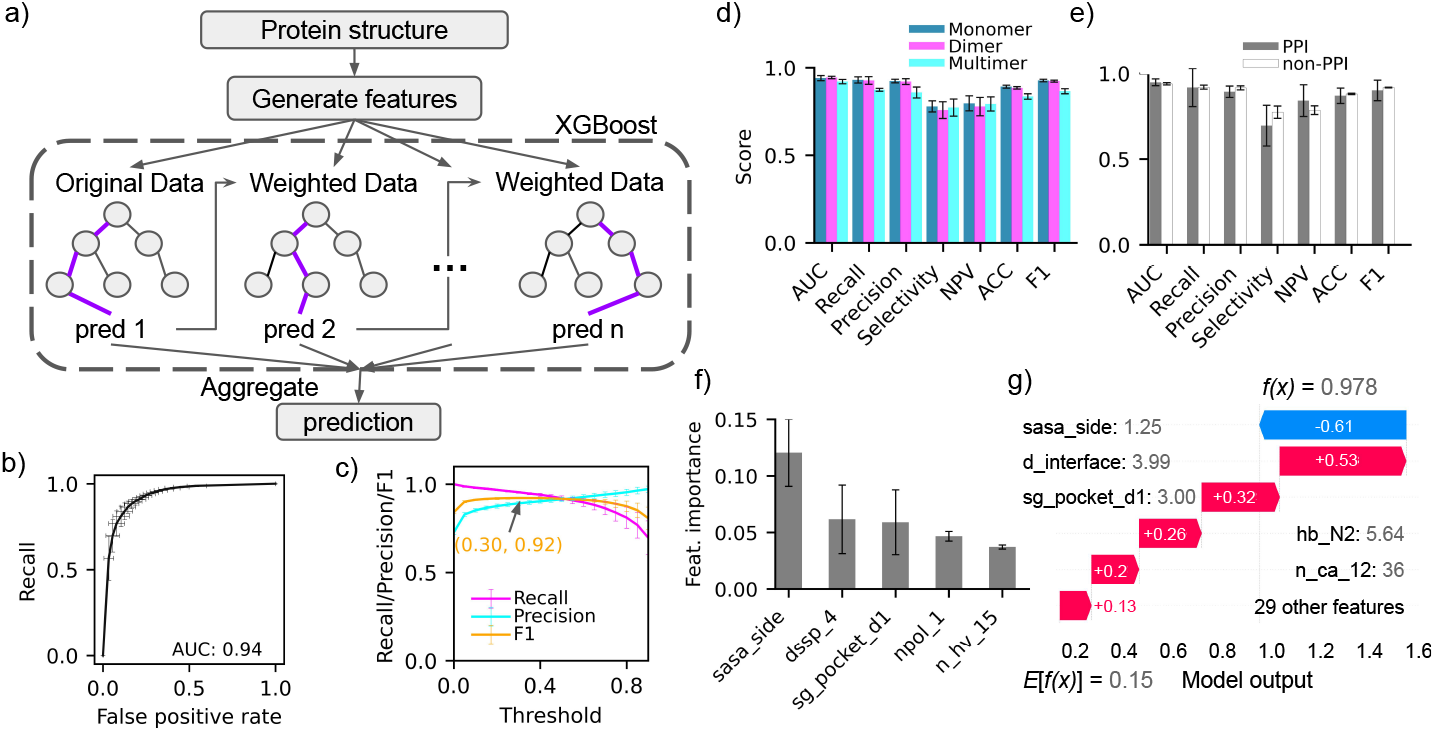
Performance of the tree-based models for predicting cysteine ligandabilities. **a)** Model workflow based on the Extreme Gradient Boosting (XGBoost) classifier. **b)** Receiver operating characteristic (ROC) curve for the XGBoost models obtained from 6 rounds of data splitting followed by training with 10-fold cross validation. The area under the curve (AUC) is 0.94. **c)** Recall/Precision/F1 score as a function of the classification threshold. The highest F1 score of 0.92 was achieved at a threshold of 0.30. **d)** Performance metrics of the ET models for cysteines in monomer, dimer, and multimer structures. **e)** Performance metrics of the ET models for cysteines at the interfaces (PPIs) or not. **f)** Permutation-based feature importance scores for the top five features: the sidechain cysteine SASA (sasa_side); secondary structure of the cysteine+4 position (dssp_4); the distance from the cysteine sulfur to the nearest pocket (sg_pocket_d1); the minimum distance to any nonpolar atom in a different residue (npol_1); and the number of heavy atoms within 15 Å of the cysteine sulfur (n_hv_15). **g)** The “Waterfall” SHAP value plot to explain the ligandability prediction for C627 in Nedd4-1’s structure (PDB: 2XBB; UniProt: P46934, C627). The five most impactful features (values are given next to the names) are shown on the top and the rest 29 features are collapsed into one and shown on the bottom. The corresponding SHAP values shown in red (positive) or blue (negative) bars accumulate to shift the expected model output *E[f(x)]* from the random guess output (0.15) to the real output (*f* (*x*) = 0.978), where *f(x)* is the model output before the logistic link function is applied.

### Model performance is unbiased with respect to protein quaternary structure and proximity to interface

It is important to verify that the model performance is unbiased with respect to the protein quaternary structures and proximity to interfaces (if any). We compared the XGBoost model performance metrics for cysteines in the monomer, dimer, and multimer structures (Fig. 2d and Supplemental Table S2. The AUCs for monomers and dimers are identical (0.94) and it is only marginally lower for multimers (0.92). While the recall or precision for monomers and dimers are also identical (0.93 or 0.92, respectively), it is only somewhat lower for multimers (0.87 and 0.86, respectively). As to non-PPI vs. PPI cysteines, the AUC, recall, and precision are nearly identical (Fig. 2e and Supplemental Table S3. These analyses demonstrate that the models are equally predictive for large protein assemblies and PPIs. The latter is desirable, as TCI discovery targeting PPIs has been very challenging. ^22^

### Cysteine ligandability is determined by a set of structural and physico-chemical features

A significant advantage of decision tree as opposed to neural network models is interpretability. The permutation feature importance scores were calculated to understand the structural and physicochemical features that determine cysteine ligandability. The feature importance score represents the decrease in the model score when a feature is randomly shuffled. ^23^ Accordingly, cysteine’s sidechain solvent-accessible surface area (sasa_side) is by far the most important feature (Fig. 2f), which is readily understood, as solvent exposure promotes cysteine reactivity due to the stabilizing solvation free energy of the anionic thiolate state. However, an earlier study found a poor correlation between the solvent accessibility and thiol reactivity. ^24^ An early bioinformatics analysis showed that cysteine is the least-exposed amino acid ^25^ and the recent constant pH MD simulations showed that many hyperreactive cysteines in kinases ^19,20^ and other proteins ^7^ are buried. We will come back to this discussion. The next four features: the secondary structure at the cysteine+4 position (dssp_4), the distance from the cysteine sulfur to the nearest pocket (sg_pocket_d1), the distance to the nearest nonpolar atom in another residue (npol_1), and the number of heavy atoms within 15 Å from the cysteine sulfur (n_hv_15), are also consistent with the intuition or knowledge from other studies. In accord with the importance score of dssp_4, the N-terminal capping (Ncap) cysteine on a helix has been suggested as highly reactive two decades ago, ^26^ which is supported by the fact that the front-pocket Ncap cysteine is the most popular site of targeted covalent inhibition among all kinases. ^19^ Similar to the BURIED term in the empirical p*K*_a_ prediction program PROPKA, ^27^ the two features npol_1 and n_hv_15 indicate how deeply the cysteine is buried, which affects both the cysteine reactivity and ligand accessibility.

Complementary to the feature importance scores, the game-theoretic SHAP (SHapley Additive exPlanations) values inform the impact of feature values on the prediction outcomes. ^28,29^ A positive or negative SHAP value increases or decreases the model output of a prediction from its expectation value estimated by randomly guessing from the features. ^29^ As an example, Fig. 2g explains the model prediction for C627 in Nedd4-1 (PDB: 2XBB) based on the SHAP values of the features. While the sasa_side is small (1.25 Å ^2^) and decreases the model output by 0.61, the other four important features, the cysteine sulfur distance to the interface (d interface, 3.99 Å), to the nearest pocket (sg_pocket_d1, 3.00 Å), and to the second nearest potential hydrogen bond donor nitrogen (hb_N2, 5.64 Å), as well as the number of C*α* atoms within 12 Å of the cysteine sulfur (n ca 12, 36) increase the model output by 0.53, 0.32, 0.26, and 0.20 respectively. Together with the 0.13 positive contribution from the rest of the features, the model output *f* (*x*) is upshifted from the expected value (*E*[*f* (*x*)]) of 0.15 to the value of 0.978, which returns a class probability score of 0.73.

### Covalent modification perturbs the cysteine structure environment

Structural perturbation by reversible ligands is a well-known phenomenon. ^30^ We hypothesized that covalent modification of a cysteine perturbs its structural environment. To test this hypothesis, we plotted the distributions of the cysteine sidechain SASA and the distance to the nearest pocket, which are important features of the tree models (Fig. 3). For the LigCys3D cysteines, the positives are separated into the modified and unmodified groups, which refer to whether the cysteine is liganded or modified in the structure or not. Note, the unmodified structures can be either apo or in complex with a reversible ligand. Interestingly, the major peak of the SASA distribution for the modified positives is at ∼20 Å ^2^, while that of the unmodified positives is at ∼5 Å ^2^, which is close to the peak of the negatives (near zero) (Fig. 3a). This analysis suggests that covalent modification perturbs the protein structure so as to increase cysteine’s solvent exposure. Furthermore, while a larger fraction of ligandable cysteines are solvent exposed as compared to the unligandable ones, a significant fraction of ligandable cysteines are actually deeply buried. The latter is consistent with the notion that cysteine is the least solvent-exposed amino acid ^25^ and our recent finding that most reactive cysteines in kinases ^8,19,20^ and other proteins are in fact buried. ^7^

**Figure 3.**
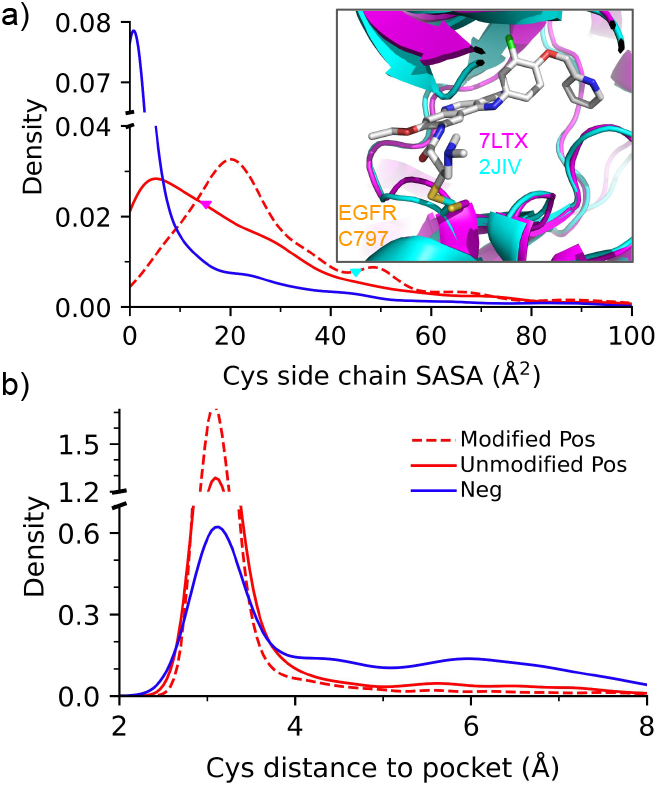
Cysteine’s conformational environment is different between the modified and unmodified structures. a) Distributions of the cysteine sidechain solvent-accessible surface area (SASA). Modified Pos (dashed red) and unmodified Pos (solid red) refer to the positive structures in which the cysteine is liganded and unliganded, respectively. Ligand was removed in the SASA calculations. The insert shows an overlay between the unmodified (PDB: 7LTX) and modified (PDB: 2JIV) X-ray structures of the EGFR kinase. The front-pocket C797 has a SASA value of 16.9 and 44.6 Å^2^ in the apo and holo states, respectively. b) Distributions of the distance from the cysteine sulfur to the nearest pocket (alpha sphere).

The distribution of the cysteine distance to the nearest pocket displays a peak near 3 Å for both modified and unmodified positives (near 3 Å, Fig. 3b); however, the modified positives have a higher peak intensity, suggesting that covalent ligand binding may slightly “pull” the cysteine towards the pocket. Interestingly, the distribution of the negatives also displays a peak near 3 Å, although with a lower peak height as compared to the positives, and importantly, the distribution has a fat tail, suggesting that many negative cysteines are far away from any pocket, as expected.

### The importance of including cysteine-unmodified structures in the training set

Given that covalent ligation perturbs the structure such that the difference in the cysteine environment between the modified positives and negatives is larger than that between the unmodified positives and negatives (Fig. 3), we hypothesized that models trained using the modified structures may give a “deceptively” higher performance than models trained with the unmodified structures. To test this, we compare the performances of the ET models trained with the modified (model 1), unmodified (model 2), and combined (model 3) structures. Confirming our hypothesis, the unseen test AUC, recall, and precision of model 1 are all above 0.95, whereas the AUC of model 2 is dropped to 0.85 and recall/precision to 0.75/0.78 (Table S6). Even though the training dataset of model 3 is the largest, 68% larger than model 1, the AUC is slightly lower at 0.94 and recall/precision is 0.89/0.93.

We compare the prediction metrics of Model 1 with the published metrics of the previous two models ^9,10^ that were trained using a smaller dataset and the cysteine-liganded structures only. Note, this comparison is for information only and needs to be taken with a grain of salt due to the difference in the training and test data. The ET metrics (AUC, recall and precision all above 0.95) surpass the feature-based SVM model (test AUC, recall and precision of 0.73, 0.62, and 0.41), ^9^ which may be attributed to the larger dataset and the use of the physio-chemical features born out of our previous mechanistic studies of cysteine reactivities and ligandabilities. ^6,19,20^ The ET model’s test AUC (0.96) is also slightly higher than the training validation AUC (0.92) of the most recent GNN model (other metrics were not reported). ^10^

Since in prospective predictions, modified structures are unavailable, we asked if the models should be trained with the unmodified structures only. To address this question, we applied the three ET models to the ligandable cysteines discovered in an early chemoproteomics experiment conducted in cell lysates. ^31^ These cysteines are not in LigCys3D, i.e., they do not have modified structures. As expected, model 1 has by far the lowest recall; however, model 3 is slightly better than model 2 in both recall and precision (Table S6). This analysis demonstrates that structure variation (including both modified and unmodified structures) further enhances the extrapolation power of the models.

### The CNN models show similar performance as the XG-Boost models

Since many of the tree model features are spatially related, we reasoned that three-dimensional convolutional neural networks (3D-CNN) may offer high performance. We adapted and modified the 3D-CNN architecture of Pafnucy which was developed for protein-ligand binding affinity predictions ^32^ and recently adapted for protein p*K*_a_ predictions. ^33^ In our modified architecture, a cubic grid of 20×20×20 Å with a resolution of 1 Å was created centering at the cysteine sulfur, and each voxel represents a nearby atom and encodes 20 features (Fig. 4a). To remove rotational variance, each cubic box was generated 20 times by randomly rotating the PDB coordinates. The input grid is processed by a block of 3D convolutional layers that have 128 filters (Fig. 4a, details see Methods). To allow comparison to the tree models, data splitting and CV were conducted in the same manner. Interestingly, the 3D-CNN gave very similar to the best tree model XGBoost, with the AUC, accuracy and precision of 0.93 ±0.04, 0.96±0.02, 0.89±0.03, respectively (Table 1). It is also noteworthy that the standard deviations in the test metrics resulting from the six data splits, training/CV, and testing are overall slightly larger than those of the XG-Boost models (Fig. 4b). Although the best average F1 score (0.92) is the same as the XGBoost models, it is achieved with a lower prediction probability threshold (0.15, Fig. 4c).

**Figure 4.**
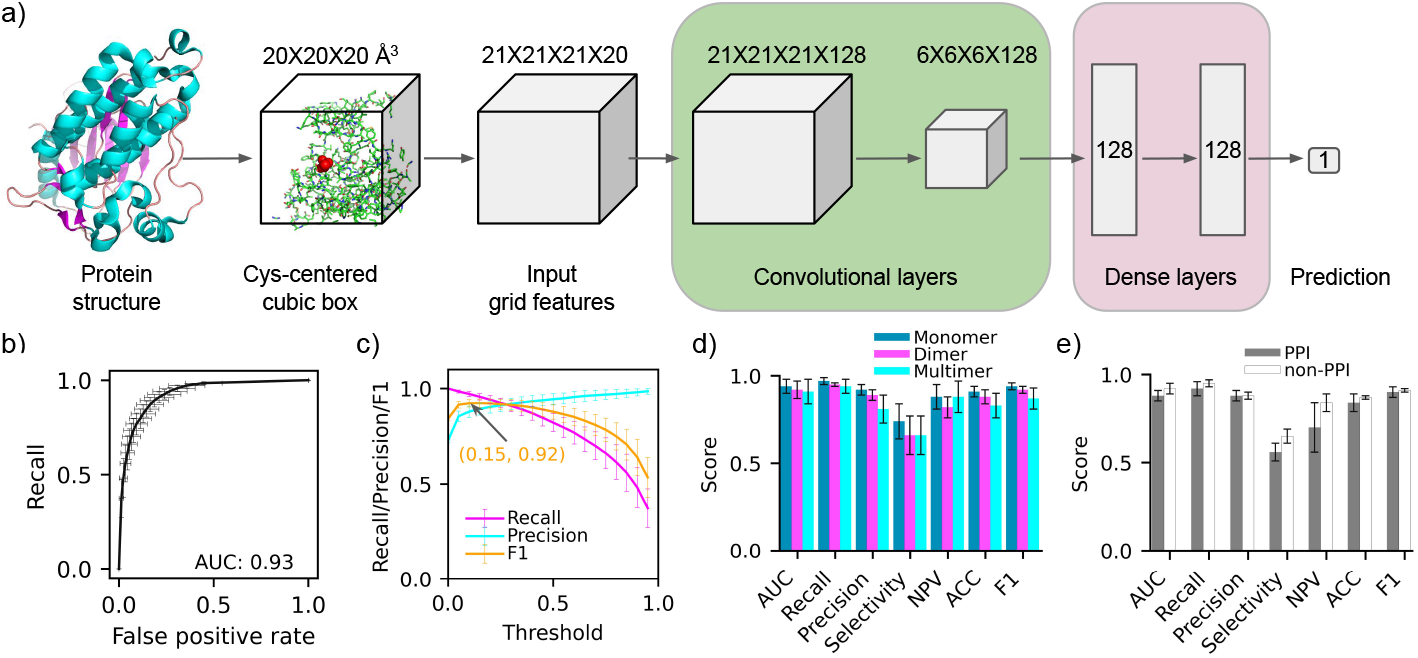
Performance of the three-dimensional convolutional neural network (CNN). **a)** Architecture of the 3D-CNN for cysteine ligandability predictions. **b)** ROC curve obtained from 6 train/test experiments. The AUC is indicated for the average curve. **c)** Recall/precision/F1 score as a function of the classification threshold. The best F1 score 0.92 is achieved at a threshold of 0.15. **d)** Comparison of the CNN performance metrics for cysteines in monomer, dimer, and multimer structures. **e)** Comparison of the CNN performance metrics for PPI and non-PPI cysteines.

We also examined the CNN performance for different protein quaternary structures and PPI vs. non-PPI cysteines in comparison to the XGBoost models (Fig. 4d and Table S4. While the AUC, recall, and precision are maintained between monomers and dimers with the XGBoost models, there is a 0.02 decrease in the average AUC or recall and 0.03 decrease in the average precision going from monomers to dimers with the CNN models. As to multimers, the average AUC or recall drop only by 0.01 relative to the dimers (smaller than the XGBoost models) but the precision drops by 0.08 (larger than the XGBoost models). This analysis suggests that the classification power of the CNN models deteriorates slightly more for dimers and multimers as compared to the XGBoost models.

The trend in the model performance differences among quaternary structures is consistent with that between the PPI and non-PPI cysteines (Fig. 4e and Table S5. While the average AUC and recall are maintained going from the non-PPI to the PPI cysteines with the XGBoost models, the respective decrease is 0.03 and 0.02 with the CNNs. As to the precision, the decrease from the non-PPI to the PPI cysteines is only 0.01 as compared to 0.03 with the XGBoost models. Interestingly, the standard deviations among the different CNN tests are doubled going from the non-PPI to the PPI cysteines, which is consistent with the XGBoost tests, although the standard deviations of the latter are overall significantly smaller. One possible reason for the performance deterioration of the CNNs for dimers and multimers is the finite-size grid which may exclude part of the chains that carries relevant information for model prediction. In contrast, the features used in the tree models cover all residues in the bioassembly regardless the distances to the cysteine of interest.

### External evaluation on the newly published ligandable cysteines captured by X-ray crystallography

To further evaluate the ET and CNN models, they were tested on a new, non-overlapping dataset of 30 unique proteins with 38 covalently liganded cysteines, for which the co-crystal structures were deposited in the PDB after the LigCys3D was constructed in Nov 2022. To demonstrate the general utility of the ML models, the AF2 structures were used for making predictions. The ET model recapitulated 35 out of 38 (a recall of 0.92), while the CNN recapitulated 34 out of 38 liganded cysteines (a recall of 0.90, see Table S7. Curiously, both the ET and CNN missed C269 in isocitrate dehydrogenases 1 (IDH1, UniProt ID O75874). Closer examination revealed that the covalently liganded IDH1 is a R132H mutant, whereas the AF2 structure used for the predictions is of the wild type (WT). Comparison of the deposited X-ray structure of the R132H mutant (PDB ID: 8HB9^34^) with the WT AF2 structure showed that the pocket near C269 is enlarged in the mutant. This difference could explain the false negative predictions. Another cysteine that both ET and CNN models missed is C412 of the YTHDF1 protein (UniProt ID Q9BYJ9). In the deposited X-ray structure (PDB ID: 7PCU. ^35^) C412 is fully exposed to solvent; however, in the AF2 structure, it is fully buried as a nearby loop (residues 95–114) is collapsed onto the pocket where the cysteine resides. These two cases show that the accuracy of the ligandability prediction is dependent on the quality of the AF2 predicted structure, which is a limitation of the structure-based ML models. Effects of mutations and loop modeling are challenging for the current AF2 model and the ligandability predictions will benefit from the continued improvement of the AlphaFold structure prediction engine.

### Prospective prediction of ligandable cysteines in the human kinome

Human kinases are important drug targets and most FDA approved covalent drugs are kinase inhibitors directed at a cysteine in the catalytic or allosteric pocket of the kinase domain. LigCys3D contains 46 unique liganded cysteines in 37 kinases, and all but two are in the kinase domain (red circles in Fig. 5). The front pocket *α*D helix, with 10 unique cysteines belonging to different kinases, is the most targeted location. This is consistent with the constant pH MD simulations which showed that the cysteine at or near the N-terminal cap of the *α*D helix is hyper-reactive due to the local hydrogen bonding and electrostatic environment. ^19^ Other popular locations for cysteine ligation are the N-terminal part of the activation loop (ALN, 8 cysteines), *α*E (5 cysteines), and the C-terminal end of *β*I (*β*IC or p-loop, 4 cysteines).

**Figure 5.**
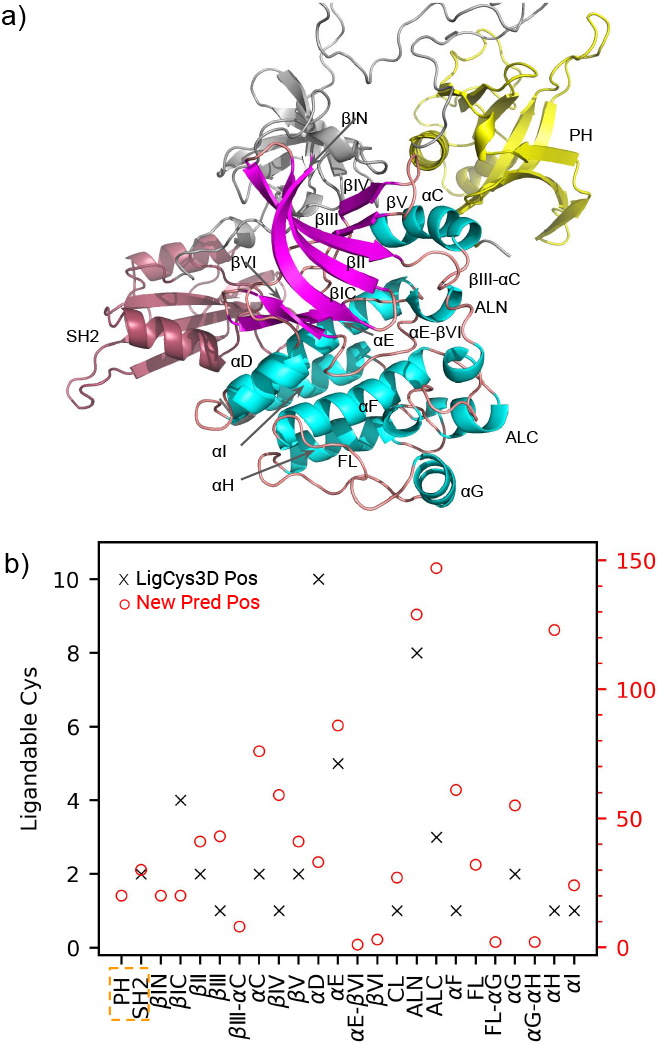
Ligandable cysteines in the kinase, PH, and SH2 domains of the human kinases. **a)** The PH (yellow), SH2 (dark red), and catalytic (magenta for *β* strands and cyan for helices) domains of a kinase are shown in a representative AF2 model structure of BTK (Uniprot ID: Q06187). The loops and regions not in the PH, SH2, or kinase domains are colored grey. The various structure elements in the kinase domain (named according to Modi and Dunbrack ^36^) as well as the PH and SH2 domains are labeled. **b)** The number of liganded cysteines in LigCys3D (grey crosses, left y axis) and the predicted ligandable cysteines for the 397 (typical) human kinases not in LigCys3D (red circles, right y axis) mapped onto the various structure elements of the kinase domain as well as the PH and SH2 domains (dashed box). The multiple sequence alignment based on the kinase domain is taken from Modi and Dunbrack. ^36^ The consensus predictions by the ET, XGBoost and LightGBM models are shown. The domain and structure element names are ordered by the sequence and only those with at least one ligandable cysteine are shown (no ligandable cysteines were predicted for the SH1 domain).

The human kinome contains 536 kinases; ^37^ however, only a small fraction (37 according to the co-crystal structures in the PDB) have been covalently drugged. Thus, we applied the ML models based on the AF2 structures to make predictions of the ligandable cysteines in the rest of the 481 kinases, accounting for not only the kinase domain but also the PH, SH1, and SH2 domains. Note, 18 kinases that do not have AF2 structure models in the EBI repository were excluded and since no ligandable cysteines were predicted in the SH1 domain, it is excluded in the discussion below.

A total of 1083 cysteines in the kinase, PH, and SH2 domains of 392 kinases were predicted to be ligandable and 89 kinases were predicted to have no ligandable cysteine. Fig. 5 shows the number of the predicted ligandable cysteines and their locations in a kinase structure according to the multiple-sequence alignment of Modi and Dunbrack. ^36^ Note, 79 predicted ligandable cysteines in 28 kinases were excluded in Fig. 5 due to the lack of sequence alignment information. Three kinase locations with most positive cysteines are ALN, the C-terminal end of the activation loop (ALC), and *α*H (Fig. 5). Other interesting locations include *α*D, *α*E, *β*IC, and *α*C. All these locations have been covalently targeted in other kinases (according to LigCys3D). Thus, our kinomewide predictions demonstrate new opportunities for covalent kinase inhibitor design but also suggest potential selectivity problems. Interestingly, the ML predictions uncovered 7 previously untargeted locations that contain predicted ligandable cysteines, including the PH domain, the N-terminal end of *β*I (*β*IN), *β*III-*α*C, *β*IV-*β*V, *α*E-*β*VI, FL-*α*G, and *α*G-*α*H (Fig. 5). Many of these locations are in the loops connecting two secondary structures.

### Development of a live online database (LigCys3D) and prediction server (DeepCys)

To assist the community of TCI discoveries, we implemented a searchable web database LigCys3D (https://ligcys.computchem.org/). Each row or entry is a ligandable cysteine in a protein, along with the following information as columns: the PDB ID, chain ID, PDB residue ID, Uniprot ID, Uniprot residue ID, bioassembly type, the calculated sidechain SASA, distance to a protein-protein interface, distance to the nearest pocket, as well as whether the cysteine is covalently modified by a ligand in the particular structure (PDB file). Any information specified in the column can be filtered and a list of matched entries are generated. The user can download the matched entries or the entire database as a CSV file. The database will be continuously updated to include the newly discovered ligandable cysteines in the PDB.

We also implemented a web server DeepCys (https://deepcys.computchem.org/) that can predict cysteine ligandability given a PDB ID, Uniprot ID, or a structure file in the PDB format. Provided a PDB ID, the web server queries the RCSB repository for the bioassembly file. Provided a Uniprot ID, the web server queries the AlphaFold Protein Structure Database of EMBL-EBI (https://alphafold.ebi.ac.uk/) for a corresponding AF2 predicted structure. Once a structure file is retrieved or provided, the server uses the models trained in this study (which will be continuously improved based on the continuously expanding training data) to make ligandability predictions for all cysteines in the protein. The results are in the format of a CSV file which contains the chain ID, residue ID, classification probability and ligandability prediction for each cysteine. The CSV file will be sent to the user-provided email address.

## CONCLUSIONS

Exploiting a newly curated database (LigCys3D) of ∼1,000 liganded cysteines in ∼800 proteins represented by ∼10,000 three-dimensional structures in the PDB, we developed the tree-based and 3D-CNN models for proteomewide cysteine ligandability predictions. In multiple unseen tests, the ET and XGBoost models gave the AUC of 94%, while the CNN models gave the AUC of 93%. Based on the AF2 predicted structures, the ET model and CNN recapitulated the newly liganded cysteines with the recall values of 92% and 90%, respectively. The fact that the tree models have orders of magnitude smaller parameter space (i.e., less than 40 structural and physico-chemical features) than CNNs reinforces the notion that the reactivity and ligandability of cysteines are largely determined by the structure environment, solvent accessibility, and potential hydrogen bonding as well as electrostatic interactions. Also, the tree models avoid the over-fitting problem facing the more sophisticated ML models that make use of very large parameter space.

Despite the promising test results, the models have several limitations. First, the training dataset is very small and under-represents transmembrane proteins, transcription factors and other non-enzymes. Second, the potential effects of mutation and post-translational modification on the structure as well as the membrane environment of transmembrane proteins are not accounted for. Third, the current AF2 engine offers very limited accuracy for predicting the loop conformations. Nonetheless, the present work represents a first step towards the ML-led integration of big genome and structure data to annotate the human proteome space for the next-generation covalent drug discoveries. To assist the community in the covalent drug design efforts, we report the predicted ligandable cysteines in 392 human kinases and their locations in the sequence-aligned kinase structure including the PH and SH2 domains. Furthermore, we disseminate a web database (https://ligcys.computchem.org/) and a web prediction server (https://deepcys.computchem.org/), both of which will be continuously updated and improved by including newly published experimental data.

## Materials and Methods

### Construction of the LigCys3D database

Two recently published databases, CovPDB ^13^ and CovalentInDB, ^10^ compiled cysteine-liganded co-crystal structures in the RCSB Protein Data bank (PDB). These two databases have overlap and together they provide 2875 cysteine-liganded co-crystal structures representing 662 liganded cysteines in 489 unique proteins. We conducted an exhaustive search in the PDB and found additionally 472 liganded cysteines in 294 unique proteins. We note, the “L-peptide linking” ^38^ cysteines that were chemically modified at locations other than the sulfur (SG) atom or simply oxidized were excluded, as well as the cysteines involved in disulfide bonds, zinc-finger coordination, or iron-sulfur clusters. Following the compilation of the cysteine-liganded structures, we used SIFTS ^14^ to annotate the liganded cysteines with UniProt accession numbers and residue IDs (https://www.uniprot.org), ^11^ which allowed us to retrieve all PDB entries associated with these cysteines. We refer to a cysteine as positive if it is liganded in any crystal structure, and the other cysteines in these structures are referred to as negatives. Note, the bioassembly structures (CIF files) were downloaded, and the coordinates of missing atoms or residues if any were added using pdbfixer (https://github.com/openmm/pdbfixer). ^39^ We refer to this dataset as LigCys3D.

### Data engineering for the ML models

To constructed a ML training set with balanced positive and negative classes and to reduce model training time, we down-sampled the number of structures in LigCys3D as follows. For each positive cysteine (based on the UniProt accession number and residue ID), all cysteine-liganded structures were included and the cysteine-unliganded structures were selected using a SASA-based protocol (see below) such that the total number of structures does not exceed 10. The cysteine-liganded and unliganded structures are referred to as the liganded and unliganded positives, respectively. For each negative cysteine, all and up to a total number of 4 structures were selected using a similar SASA-based protocol (see below). To maximize structural variation, the unliganded positive structures were put into four bins based on the cysteine sidechain solvent accessible surface area (SASA) values, and one structure was randomly picked from each bin. Similarly, the structures representing a negative cysteine were put into ten bins based on the SASA values, and one structure was randomly picked from each bin. Subsequently, a training dataset (down-sampled from LigCys3D) was constructed, comprising 9,992 positive (1,133 unique positive cysteines in 9,992 structures) and 10,267 negative (3,084 unique negative cysteines in 10,267 structures) entries. We will use this dataset for model training and testing.

### Feature engineering for the tree models

Features are critical for the performance of tree-based models. We conceived a set of structural and physico-chemical features based on our findings from the constant pH molecular dynamics (MD) analysis of cysteine reactivities and ligandabilities in a large number of kinases ^6,8,19,20^ and other enzymes. ^7^ In total, eights types of features were calculated based on the input structure, including solvent accessibility (proximity to hydrophobic residues and the cysteine SASA calculated with NACCESS ^40^); distance to pockets (defined by fpocket ^41^); potential hydrogen bonding; electrostatic interactions; secondary structures (calculated with Biopython ^42^); residue flexibility (calculated with PredyFlexy ^43^); distance to protein-protein/nucleic acid interface; and presence or absence of ligand binding. A detailed list of the features that were tested is given in Supplemental Methods. After removing highly correlated features, 37 features were left (see Supplemental Methods).

### Data splitting and training of the tree models

PyCaret ^21^ was used for building tree-based classifiers. We manually separated 10% of the data as the unseen test set and 90% as training set (see below). The training set was used for the 10-fold cross validations (CVs). To ensure that the training and testing sets do not contain structures representing the same cysteine, we first grouped the structures according to the UniProt residue IDs and then performed the training-test split by the UniProt residue IDs. In the cross validation, the groupKfold method was used to avoid placing (modified or unmodified) structures associated the same cysteine in different folds. Note, since ligandability of a cysteine is mainly determined by its local conformational environment, ^6–8^ amino acid sequence based clustering such as that employed in the previous work ^9,10^ was not used for data splitting.

Multicollinearity was removed with a threshold of 0.9. This leads to a total of 37 features (see above). Categorical features were one-hot encoded. Model training used the binary cross-entropy as loss function and default hyperparameters. The default scikit-learn search library was used to search the hyper-parameters, which were tuned using the tune model function in PyCaret 5000 times by optimizing the F1 score across all validation folds. Following tuning, the best hyperparameters were used to train the entire training set, and the final model was saved for predictions on the unseen test set or the ABPP dataset. Feature importance scores were generated using the evaluate model function. To generate statistics for model evaluation, the above process was repeated 6 times, and the average and standard deviation of the model performance metrics were calculated.

### Training of convolutional neural networks (CNNs)

The test-train splitting and 10-fold cross validation were performed in the same manner as the tree models. The 3D-CNN architecture was adapted and modified from the Pafnucy model, ^32^ which was recently adapted for protein p*K*_a_ predictions. ^33^ The input of the CNN represents a 3D image of the protein with 20 color channels. Specifically, a 20-Å 3D grid centered at the SG atom of the cysteine of interest was created. The protein heavy atoms were mapped to the grid with a 1-Å resolution, and each grid point was encoded with 20 features (the default is zero if no atoms): one-hot encoding of 5 atom types C, N, O, S, and others; 1 integer (1/2/3) for atom hybridization; 1 integer for the number of bonded heavy atoms; 1 integer for the number of bonded hetero atoms; one-hot encoding (5 in total) of the SMARTS patterns ^44^ hydrophobic, aromatic, acceptor, donor, and ring; 1 float for grid charge; one-hot encoding of 6 residue types Asp/Glu, Lys/Arg, His, Cys, Asn/Gln/Trp/Tyr/Ser/Thr, and others. Each cubic box was generated 20 times by rotating the coordinates in the PDB structure to remove rotational variance.

Keras 3^45^ was used to build the CNN. The CNN model contains two Conv3D layers and each Conv3D layer has 128 filters, kernel size 5, activation function relu, and ‘same’ padding, followed by a pool size 2 MaxPool3D layer and a BatchNormalization layer. Next, a GlobalAveragePooling3D layer is added to do global pooling and then the data is flatterned by a 128 units Dense layer with relu activation, normalized by a BatchNormalization layer, and filtered by a 0.5 ratio Dropout layer. Finally, a Dense layer of 1 unit and sigmoid activation function is used to generate a binary classification result. Batch size is set to 32 and binary cross-entropy is used as loss function. 50 epochs of training in Adam optimizer is used, with the learning rate of 0.0001 and early stopping if the validation loss plateaus in 5 epochs. The model with the lowest loss in the validation set is saved for the tests on the unseen LigCys3D data and the new AF2 data. In these tests, we used the voting result based on the predictions by the 10 saved models from CVs. The voting threshold was determined by the average F1 score on the test set across 6 train/CV:test splitting experiments. For the unseen and external testing, the predictions were determined by the majority voting.

### External validation on the newly liganded cysteines

Structure files that were deposited in the PDB between 11/29/2022 and 10/11/2023 and have publications were retrieved and analyzed as to 1) whether the cysteines are covalently modified as defined above; 2) whether the corresponding protein is present in LigCys3D; and 3) whether an AF2 structure corresponding to the Uniprot accession number is available in the EMBL-EBI AlphaFold repository (https://alphafold.ebi.ac.uk/). The AF2 structures were subsequently used in the ligandability predictions.

### Prediction of ligandable cysteines in the human kinome

We collected the human kinase list from KinMap. ^37^ For those with AF2 predicted structures, ligandability predictions were made using three tree methods (ET, XGBoost, and Light-GBM) and the consensus scheme. The kinase domain information was extracted from the Uniprot *Family & Domains* section and if the domain name starts with text *Kinase domain*, the corresponding residue range is considered as a kinase domain for that Uniprot ID. We also used the KinCoRe alignment file ^36^ downloaded from the website http://dunbrack.fccc.edu/kincore/biojs to assign the structure location for each cysteine in the kinase domain. For cysteines in the PH, SH2, and SH1 domains, if the Uniprot domain name starts with them, the original domain names are renamed as the short forms so that multiple SH2 domains are combined. For those cysteines that are in the kinase domain but are not found in KinCore, the domain name is just *Kinase domain* but they are not shown in Fig. 5. Cysteines not in the kinase, PH, SH1 and SH2 domains are not discussed in this study. For the location names in KinCore, *A* was replaced with *α* to indicate *α*-helix, *B* was replaced with *β* to indicated *β*-sheet, and an Arabic number was replaced by a Roman number. These changes were made to be more consistent with the literature.

### Calculation of model performance metrics

Given a confusion matrix comprised of the number of true positives (TPs), true negatives (TNs), false positives (FPs), and false negatives (FNs), the model performance metrics, recall (or true positive rate TPR), precision, specificity, negative predictive value (NPV), accuracy (ACC), and F1 score are defined as follows.

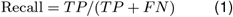

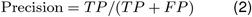

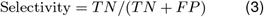

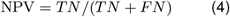

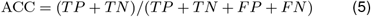

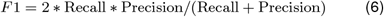

AUC is calculated by integrating the area under the ROC (receiver operating characteristic) curve, which consists of the recall and false positive rate (1 - selectivity) at all possible classification threshold values.

## Supporting Information Available

Supporting Information contains supplemental methods, tables, and figures. The predicted ligandable cysteines in 392 kinases and their locations are provided as a downloadable file (Kinase Ligandable Cys.xlsx).

## Data Availability

All training and testing data as well as the ML models are downloadable from https://github.com/JanaShenLab/DeepCys. The training dataset Lig-Cys3D is also provided on the web as a searchable database (https://ligcys.computchem.org/). The tree models and CNNs are also provided as a web server DeepCys (https://deepcys.computchem.org/).

## Acknowledgement

We acknowledge financial support by the National Cancer Institute (R01CA256557).

## TOC Graphic

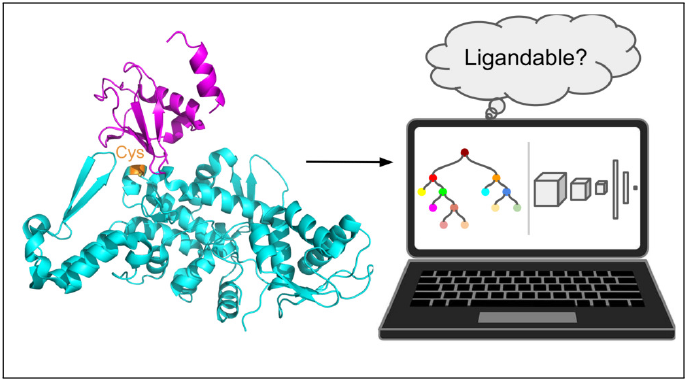

## Notes

https://github.com/JanaShenLab/DeepCys

### Competing Interest Statement

J.S. is a scientific advisory board member of Matchpoint Therapeutics.

### Summary of Updates

The Results and Discussion has been revised. New data and analysis are added. Web servers are added for the database and prediction engine.

