## Supplemental tables for "Machine Learning Models to Interrogate Proteomewide Covalent Ligandabilities Directed at Cysteines"

#### Supplemental Methods

**Construction of the LigCys3D database.** The following "L-peptide linking" cysteines were excluded from the liganded cysteine data set: CSD, OCS, CSX, CSO, SNC, CSP, CZZ, CSU, 2CO, CSS, SMC, CY3, BB9, CG6, CZ2, 03Y, CMT, CSZ, DCY, CSB, CMH, BB7, BB6, 00C, XCN.

**Feature engineering for the tree models.** We first tested the following features for the tree models.

1. Solvent accessibility:

   sasa_side and sasa_mainchain: solvent accessible surface area (SASA) of the Cys sidechain and mainchain calculated by Naccess V2.1.1 (`www.bioinf.manchester.ac.uk/naccess`) based on the Lee& Richards method[S1] with a probe radius of 1.4 Å;

   npol_1 and npol_2: distances from the Cys SG atom to the nearest and second nearest nonpolar atoms in other residues;

   n_ca_6/9/12/15: number of C$\alpha$ atoms within 6/9/12/15 Å of the Cys SG atom;

   n_hv_6/9/12/15: number of heavy atoms within 6/9/12/15 Å of the Cys SG atom. pol_1/2: distance from SG to the first and second nearest polar atom in another residue;

2. Potential hydrogen bonding:

   hb_N1 and hb_N2: distances from the Cys SG atom to the first and second nearest sidechain nitrogen atoms, including ND1/NE2 in His, ND2 in Asn, NE2 in Gln, and NE1 in Trp;

   hb_O1 and hb_O2: distances from the Cys SG atom to the first and second nearest sidechain oxygen atoms, including OG in Ser, OG1 in Thr, and OH in Tyr; bb_N1 and bb_N2: distances from the Cys SG atom to the first and second nearest backbone amide nitrogen atoms of other residues.

3. Electrostatic interactions:

   pos_N1 and pos_N2: distances from the Cys SG atom to the first and second nearest nitrogen atoms of positively charged sidechains, including NZ in Lys and NE/NH1/NH2 in Arg;

   neg_O1 and neg_O2: distance from the Cys SG atom to the first and second nearest oxygen atoms of negatively charged sidechains, including OE1/OE2 in Glu and OD1/OD2 in Asp.

4. Residue flexibility:

   rmsf/bvalue: Cys flexibility RMSF and B-factor calculated using PredyFlexy;[S2]

5. Distance to another chain:

   d_interface: minimum distance between the Cys C$\alpha$ atom and any heavy atom of another chain in the biological assembly.

6. Proximity to a pocket:

   ca/sg_pocket_d1/2: distances from the Cys C$\alpha$ or SG atom to the alpha sphere in the nearest and second nearest pockets identified by fpocket;[S3]

7. Secondary structures:

   dssp_1, dssp_2, dssp_4: secondary structure of the Cys, Cys+2, and Cys+4 positions calculated using Biopython[S4] based on the definition of DSSP;[S5]

8. Ligand perturbation:

   ligand_weight: largest molecular weight of any ligand bound to the chain.

Unless otherwise noted, the features were calculated using the in-house Python scripts. The feature value was set to 999, if it is not applicable, e.g., in apo structures, ligand weight is not applicable.

After removing highly correlated features (using the option 'remove multicolinearity' in PyCaret[S6] with a threshold of 0.9), the following 37 features were left: sasa_side; sasa_mainchain; n_ca_6, n_ca_9, n_ca_12; n_hv_6, n_hv_9, n_hv_15; sg_pocket_d1, sg_pocket_d2, ca_pocket_d1, ca_pocket_d2; bb_N1, bb_N2; hb_h_1; hb_n_2; hb_O1, npol_1, npol_2; neg_O1; pos_N1, pos_N2; n_dssp_L, n_dssp_H, n_dssp_B, n2_dssp_L, n2_dssp_H, n2_dssp_B, n2_dssp_N, n4_dssp_L, n4_dssp_H, n4_dssp_B, n4_dssp_N, d_interface; rmsf, bvalue; largest_ligand_weight.

### Supplemental Tables

Table S1: Performance metrics of the Extra Trees model with 30 random train-test splittings

| Metrics | CV | Test |
|---|---|---|
| AUC | 0.89±0.00 | 0.94±0.01 |
| Recall | 0.81±0.01 | 0.90±0.02 |
| Prec | 0.77±0.01 | 0.93±0.01 |
| Select | 0.82±0.01 | 0.82±0.02 |
| NPV | 0.85±0.00 | 0.75±0.03 |
| ACC | 0.81±0.00 | 0.88±0.01 |
| F1 | 0.79±0.00 | 0.91±0.01 |

Table S2: Performance metrics of the XGBoost models for different protein quaternary structures[a]

| Metrics | Monomers | Dimers | Multimers |
|---|---|---|---|
| AUC | 0.94±0.01 | 0.94±0.01 | 0.92±0.01 |
| Recall | 0.93±0.02 | 0.93±0.02 | 0.87±0.01 |
| Prec | 0.92±0.01 | 0.92±0.02 | 0.86±0.03 |
| Select | 0.78±0.03 | 0.76±0.05 | 0.77±0.05 |
| NPV | 0.80±0.04 | 0.78±0.05 | 0.79±0.04 |
| ACC | 0.89±0.01 | 0.89±0.01 | 0.84±0.02 |
| F1 | 0.93±0.01 | 0.92±0.01 | 0.87±0.01 |

[a]The calculated metrics are the average and standard deviation from 6 train, CV, and test experiments (also given in Fig. 2d).

Table S3: Performance metrics of the XGBoost models for non-PPI and PPI cysteines

| | Non-PPI | PPI |
|---|---|---|
| AUC | 0.94±0.01 | 0.95±0.02 |
| Recall | 0.92±0.01 | 0.92±0.01 |
| Prec | 0.92±0.01 | 0.89±0.03 |
| Select | 0.77±0.04 | 0.70±0.12 |
| NPV | 0.79±0.03 | 0.84±0.09 |
| ACC | 0.88±0.00 | 0.87±0.04 |
| F1 | 0.92±0.00 | 0.90±0.06 |

[a]The calculated metrics are the average and standard deviation from 6 train, CV, and test experiments (also given in Fig. 2e).

Table S4: Performance metrics of the CNN models for different protein quaternary structures[a]

| | Monomers | Dimers | Multimers |
|---|---|---|---|
| AUC | 0.94±0.04 | 0.92±0.05 | 0.91±0.07 |
| Recall | 0.97±0.02 | 0.95±0.01 | 0.94±0.04 |
| Prec | 0.92±0.03 | 0.89±0.03 | 0.81±0.08 |
| Select | 0.74±0.10 | 0.66±0.11 | 0.66±0.11 |
| NPV | 0.88±0.07 | 0.82±0.06 | 0.88±0.09 |
| ACC | 0.91±0.03 | 0.88±0.04 | 0.83±0.07 |
| F1 | 0.94±0.02 | 0.92±0.02 | 0.87±0.06 |

[a]The calculated metrics are the average and standard deviation from 6 train, CV, and test experiments (also given in Fig. 3d).

Table S5: Performance metrics of the CNN models for the non-PPI and PPI cysteines[a]

| | Non-PPI | PPI |
|---|---|---|
| AUC | 0.93±0.04 | 0.90±0.08 |
| Recall | 0.96±0.02 | 0.94±0.05 |
| Prec | 0.89±0.03 | 0.88±0.06 |
| Select | 0.70±0.10 | 0.59±0.13 |
| NPV | 0.86±0.06 | 0.78±0.15 |
| ACC | 0.89±0.04 | 0.86±0.07 |
| F1 | 0.92±0.02 | 0.91±0.05 |

[a]The calculated metrics are the average and standard deviation from 6 train, CV, and test experiments (also given in Fig. 2e).

Table S6: Impact of training with unmodified structures on the ET model predictions[a]

| Model[b] | Model 1 | Model 2 | Model 3 |
|---|---|---|---|
| Structures | Modified | Unmodified | Combined |
| Pos:Neg[c] | 5931:5931 | 4061:4061 | 9992:10267 |
| AUC | **0.96**±**0.00** | 0.85±0.02 | 0.94±0.00 |
| Recall | **0.95**±**0.01** | 0.75±0.03 | 0.89±0.02 |
| Prec | **0.96**±**0.01** | 0.78±0.03 | 0.93±0.01 |
| F1 | **0.95**±**0.01** | 0.77±0.03 | 0.91±0.01 |
| | **Chemical proteomics (179:429)**[c,d] | | |
| AUC | 0.71±0.01 | 0.72±0.01 | **0.72**±**0.01** |
| Recall | 0.67±0.02 | 0.81±0.02 | **0.82**±**0.02** |
| Prec | 0.72±0.00 | 0.67±0.01 | **0.69**±**0.00** |
| F1 | 0.70±0.01 | 0.72±0.01 | **0.73**±**0.01** |

[a]Average and standard deviation of the metrics from the six model predictions are given. The metrics of the best model are highlighted in bold font. [b]Model 1, Model 2, and Model 3 refer to the ET models trained with the cysteine-liganded, cysteine-unliganded, and combined structures, respectively. [c]The number of positives and negatives in the entire dataset (training, CV, and unseen test). [d] This dataset is taken from an early study based on the the isotopic tandem orthogonal proteolysis (isoTOP) activity based protein profiling (ABPP) experiments in cell lysates.[S7] Note, the predictions are based on unmodified X-ray structures are used. The proteins do not overlap with those in LigCys3D. The precision and F1 score are proportionally adjusted based on the null model values of 0.29 and 0.37, respectively.

Table S7: Model evaluation on the newly published covalently-modified cysteines

| Uniprot ID | Gene | Uniprot Resid | ET | CNN |
|---|---|---|---|---|
| O14508 | SOCS2_HUMAN | 111 | True | True |
| P53539 | FOSB_HUMAN | 172 | True | True |
| Q9NYL2 | M3K20_HUMAN | 22 | True | True |
| P11392 | PHEA_PORPP | 82 | True | True |
| P11392 | | 139 | True | True |
| P11393 | PHEB_PORPP | 50 | True | True |
| P11393 | | 61 | True | True |
| P11393 | | 82 | True | True |
| P11393 | | 158 | True | True |
| P37330 | MASZ_ECOLI | 617 | True | True |
| P45984 | MK09_HUMAN | 116 | True | True |
| P02699 | OPSD_BOVIN | 322 | True | True |
| P35991 | BTK_MOUSE | 481 | True | True |
| Q5W0Q7 | USPL1_HUMAN | 236 | True | True |
| Q9ZI86 | LNT_PSEAE | 113 | True | True |
| Q9ZI86 | | 382 | True | True |
| O75874 | IDHC_HUMAN | 269 | False | False |
| E9K9Z1 | GCCF_LACPN | 64 | True | False |
| Q8IDK7 | Q8IDK7_PLAF7 | 495 | False | True |
| Q9RZA4 | BPHY_DEIRA | 24 | True | True |
| Q8ZNR3 | SOPA_SALTY | 753 | True | True |
| D6YWY5 | D6YWY5_WADCW | 313 | True | True |
| P0ADG7 | IMDH_ECOLI | 305 | True | True |
| Q06830 | PRDX1_HUMAN | 173 | True | True |
| Q9BQF6 | SENP7_HUMAN | 992 | True | True |
| P33767 | OSTB_YEAST | 400 | True | True |
| A0A067XP79 | A0A067XP79_9CRYP | 67 | True | False |
| A0A067XP89 | A0A067XP89_9CRYP | 50 | True | True |
| A0A067XP89 | | 61 | True | True |
| A0A067XP89 | | 82 | True | True |
| A0A067XP89 | | 158 | True | True |
| Q9BYJ9 | YTHD1_HUMAN | 412 | False | False |
| G2LDR3 | G2LDR3_CHLTF | 21 | True | True |
| A0A0H3JDV8 | A0A0H3JDV8_ECO57 | 753 | True | True |
| P93025 | PHOT2_ARATH | 426 | True | True |
| A2FQ38 | A2FQ38_TRIV3 | 53 | True | True |
| Q9NXF7 | DCA16_HUMAN | 58 | True | True |
| A0A045JB88 | A0A045JB88_MYCTX | 143 | True | True |

(S6) Ali, M. PyCaret: An Open Source, Low-Code Machine Learning Library in Python. 2020.

(S7) Backus, K. M.; Correia, B. E.; Lum, K. M.; Forli, S.; Horning, B. D.; González-Páez, G. E.; Chatterjee, S.; Lanning, B. R.; Teijaro, J. R.; Olson, A. J.; Wolan, D. W.; Cravatt, B. F. Proteome-Wide Covalent Ligand Discovery in Native Biological Systems. *Nature* **2016**, *534*, 570–574.
